# The SERCA residue Glu340 mediates inter-domain communication that guides Ca^2+^ transport

**DOI:** 10.1101/2020.06.22.165365

**Authors:** Maxwell M. G. Geurts, Johannes D. Clausen, Bertrand Arnou, Cedric Montigny, Guillaume Lenoir, Robin A. Corey, Christine Jaxel, Jesper V. Møller, Poul Nissen, Jens Peter Andersen, Marc le Maire, Maike Bublitz

## Abstract

The sarco(endo)plasmic reticulum Ca^2+^-ATPase (SERCA) is a P-type ATPase that transports Ca^2+^ from the cytosol into the SR/ER lumen, driven by ATP. This primary transport activity depends on tight coupling between movements of the transmembrane helices forming the two Ca^2+^ binding sites and of the cytosolic headpiece mediating ATP hydrolysis. We have addressed the molecular basis for this intramolecular communication by analyzing the structure and functional properties of the SERCA mutant E340A. The mutated Glu340 residue is strictly conserved among the P-type ATPase family of membrane transporters and is located at a seemingly strategic position at the interface between the phosphorylation domain and the cytosolic ends of five out of SERCA’s ten transmembrane helices. The mutant displays a marked slowing of the Ca^2+^-binding kinetics, and its crystal structure in the presence of Ca^2+^ and ATP analogue reveals a rotated headpiece, altered connectivity between the cytosolic domains and altered hydrogen bonding pattern around residue 340. Supported by molecular dynamics simulations, we conclude that the E340A mutation causes a stabilization of the Ca^2+^ sites in a more occluded state, hence displaying slowed dynamics. This finding underpins a crucial role of Glu340 in inter-domain communication between the headpiece and the Ca^2+^-binding transmembrane region.

## Introduction

The Ca^2+^-ATPase of sarco(endo)plasmic reticulum (SERCA) is an ion-translocating ATPase belonging to the P-type family of membrane transporters. It pumps cytosolic Ca^2+^ ions across the sarco- or endoplasmic reticulum membrane at the expense of ATP, a vital function in all living cells, particularly in the context of muscle contraction, Ca^2+^ signaling and cell survival (1).

The molecular Ca^2+^ transport mechanism of SERCA is based on a cyclic transition through different conformations of the 110 kDa membrane protein, allowing alternating access to the Ca^2+^ binding sites from the cytosol and the SR/ER lumen. Binding and hydrolysis of ATP is mediated by SERCA’s cytosolic ‘headpiece’ which consists of three roughly globular domains, the nucleotide-binding (N), phosphorylation (P), and actuator (A) domain, while the Ca^2+^ binding sites are located in the transmembrane (TM) part of the protein, which consists of ten α-helices (M1-M10) (Fig 1A). ATP binding and the subsequent transient phosphorylation of a conserved aspartate residue are coupled to a sequence of binding of two Ca^2+^ ions from the cytosol, followed by occlusion (a state where the ions are shielded from access to either side of the membrane), and luminal release of these ions. This is associated with conformational transitions between so-called *E*1, *E*1P, *E*2P and *E*2 forms (Fig 1B, ‘P’ representing phosphorylation and brackets representing ion occlusion)(1, 2).

**Figure 1.**
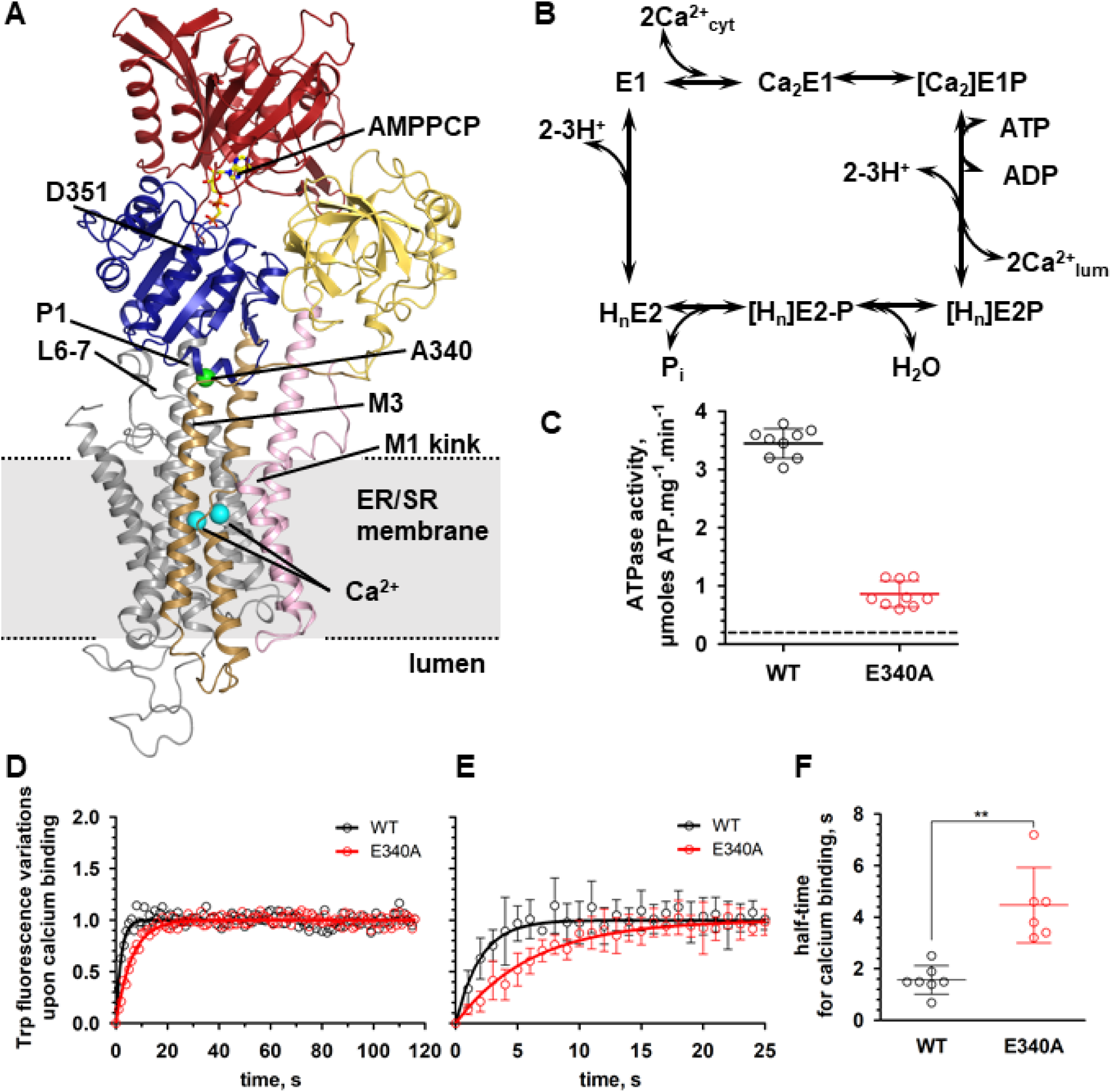
SERCA reaction scheme, structure and activity measurements. A Crystal structure of SERCA E340A shown as cartoon. Nucleotide-binding (N) domain is red, actuator (A-) domain yellow, phosphorylation (P-) domain blue, M1-2 pink, M3-4 brown, M5-10 grey. AMPPCP is shown as ball-and-stick, Ca^2+^ ions as cyan spheres. Residue 340 is indicated by a green sphere. B Schematic reaction cycle of SERCA. C Steady-state ATPase activity of purified WT SERCA (black) or E340A mutant (red) in an enzyme coupled assay. WT: 3.45 +/- 0.26 (n=9); E340A: 0.86 +/- 0.22 (n=9); background (dashes): 0.18 +/- 0.06 (n=18). 3 independent experiments were done from 2 different batches of purified protein. D Time course of Ca^2+^ binding transition determined by tryptophan fluorescence. Each time point represents the average of several measurements (n=7 for WT and n=6 for E340A from two independent experiments). E Magnified view of the early time points in D. Error bars correspond to the standard deviation from the mean (n=7 for WT and n=6 for E340A). F Half-time for Ca^2+^ binding for WT SERCA and E340A. t_1/2_ were determined on each individual experiment. t_1/2_ are 1.6 +/- 0.6 s for WT and 4.5 +/- 1.5 s for E340A with p=0.0023 (one-tailed paired “t” test).

A number of crystal structures have shed light on the nature of the conformational changes associated with Ca^2+^ transport (reviewed in (3–5)). There are two Ca^2+^-binding sites within the TM domain of SERCA, denoted sites I and II based on a proven sequential order of Ca^2+^ binding (6). The Ca^2+^ coordination in site I is mediated by residues in helices M5, M6, and M8, and in site II by residues in M4 and M6. Ca^2+^ binding from the cytosol to SERCA involves the characteristically kinked M1 helix (Fig 1A): a large vectorial movement of M1 toward the luminal side of the membrane mediates Ca^2+^ access from the cytosolic side, whereas Ca^2+^ occlusion after binding depends on an M1 movement in the opposite direction, allowing the formation of a hydrophobic cluster around the kink that shields the Ca^2+^ sites. This goes along with a smaller shift of M3 relative to M5 (7, 8). The transition from the *E*1P to the *E*2P state is associated with a large rotation of the A domain, which causes a distortion of the coordination geometry at the high-affinity Ca^2+^-binding sites and a concomitant loss of Ca^2+^ affinity, along with the formation of a luminal exit pathway for Ca^2+^ release (9).

Both the P and the A domain of the cytosolic headpiece are in contact with the ten-transmembrane helical domain, albeit by different structural elements: the A domain is connected via three loops (A-M1, M2-A and A-M3) that are thought to relay the movements of the A domain to the M1-M3 helices. The P domain however, is structurally very tightly integrated into the cytosolic ends of M4 and M5. M4 is linked to the so-called P1 helix (Pro337-Cys344), a short α-helix that runs roughly parallel to the membrane surface, at the membrane-facing side of the P domain. In such a location, P1 may be a key element of inter-domain communication: it connects directly to a β-strand in the P domain ending with the phosphorylated aspartate residue, and it makes contact to the cytosolic end of M3 and to the loop between M6 and M7 (L6-7), which has been shown to play an important role in SERCA catalysis (10–16). (Fig 1A).

To understand the Ca^2+^ transport mechanism it is mandatory to obtain information on the structural relations of residues critical in mediating the communication between the membranous Ca^2+^ binding sites and the cytosolic phosphorylation site. Glu340 is a centrally positioned residue in the P1 helix at the interface between the phosphorylation domain and the cytosolic ends of M3-M7 (Fig 1A). It is almost universally conserved throughout the large P-type ATPase superfamily. With the exception of polyamine transporting pumps and some bacterial representatives that mostly have a glutamine or asparagine residue at this position, all animal P-type ATPases known to pump ions or lipids strictly possess glutamate (Table S1) (17).

Mutation of Glu340 seems to affect the partial reactions involving cytoplasmic Ca^2+^ interactions (16), thus raising the question whether this residue plays an important role in linking the P and the TM domains. Structural information supporting such a role has, however, been lacking.

In general, structural information on SERCA mutants is relatively scarce with only four published crystal structures to date (18–20). The main reason for this shortage is that purification and crystallization of recombinantly produced SERCA is challenging, with low protein yields and poor stability in the absence of native lipids compared to the native enzyme purified from skeletal muscle.

In this study, we have determined the crystal structure of yeast-expressed rabbit SERCA1a mutant E340A at 3.2 Å in the Ca_2_*E*1 form with bound ATP analogue (AMPPCP), and we examined its functional properties, including ATPase activity and Ca^2+^ binding kinetics. This is notably the first determination of the structure of a SERCA mutant being catalytically active, *i*.*e*. capable of completing a full catalytic cycle, albeit with altered kinetics. Furthermore, molecular dynamics (MD) simulations of both wild-type (WT) and E340A structures embedded in a lipid bilayer supported our conclusions derived from structural and functional studies. Our data link the structure to Glu340’s functional importance and provide insight into the inter-domain communication that guides Ca^2+^ transport by SERCA.

## Results

### SERCA E340A has a lower ATPase activity and slower Ca^2+^ binding kinetics than WT

The effect of the E340A mutation on yeast-expressed SERCA function was addressed by using an enzyme-coupled assay to estimate the specific activity of the E340A mutant in detergent. We observed that the ATPase activity of the E340A mutant is about 25 % of the WT (0.86 vs. 3.45 µmoles ATP/mg/min, n=9) (Fig 1C, FigS1), *i*.*e*. a marked slowing of the enzyme cycle.

In order to assess whether this overall deceleration is due to a reduced rate of Ca^2+^ binding to SERCA E340A, we measured the time course of the Ca^2+^ binding transition, by recording changes in intrinsic tryptophan fluorescence upon addition of Ca^2+^ to the *E*2 state of the protein. The *E*2 to Ca_2_*E*1 transition of E340A is significantly slower than that of the WT, with an increase of t_1/2_ of about 3-fold (Fig 1D-F). This indicates that the effect of the E340A mutation on the overall turnover of SERCA is caused by a delay in the conformational change associated with Ca^2+^ binding, i.e. the H_n_*E*2 -> *E*1 -> Ca_2_*E*1 transition.

### The mutation E340A causes global changes in SERCA’s domain arrangement

Our crystal structure of SERCA E340A at 3.2 Å resolution (Fig 1A, Table S1) allows for a detailed structural comparison with the WT structure. When superposed, the WT and E340A structures deviate by a substantial RMSD of 2.3 Å over all main chain atoms. Nevertheless, when superposed on the C-terminal transmembrane helix bundle M5-M10 (residues 750-994, main chain RMSD= 0.49 Å), the E340A mutant and WT have an almost identical arrangement of the remaining transmembrane helices M1-M4 (main chain RMSD = 0.56 Å), with a small 1.6 Å bend at the cytosolic end of M3 in E340A (Fig 2A), and a slightly more kinked helix M1 (105° in E340A vs. 109° in WT). In line with this, the Ca^2+^ binding sites look identical in WT and E340A.

**Figure 2.**
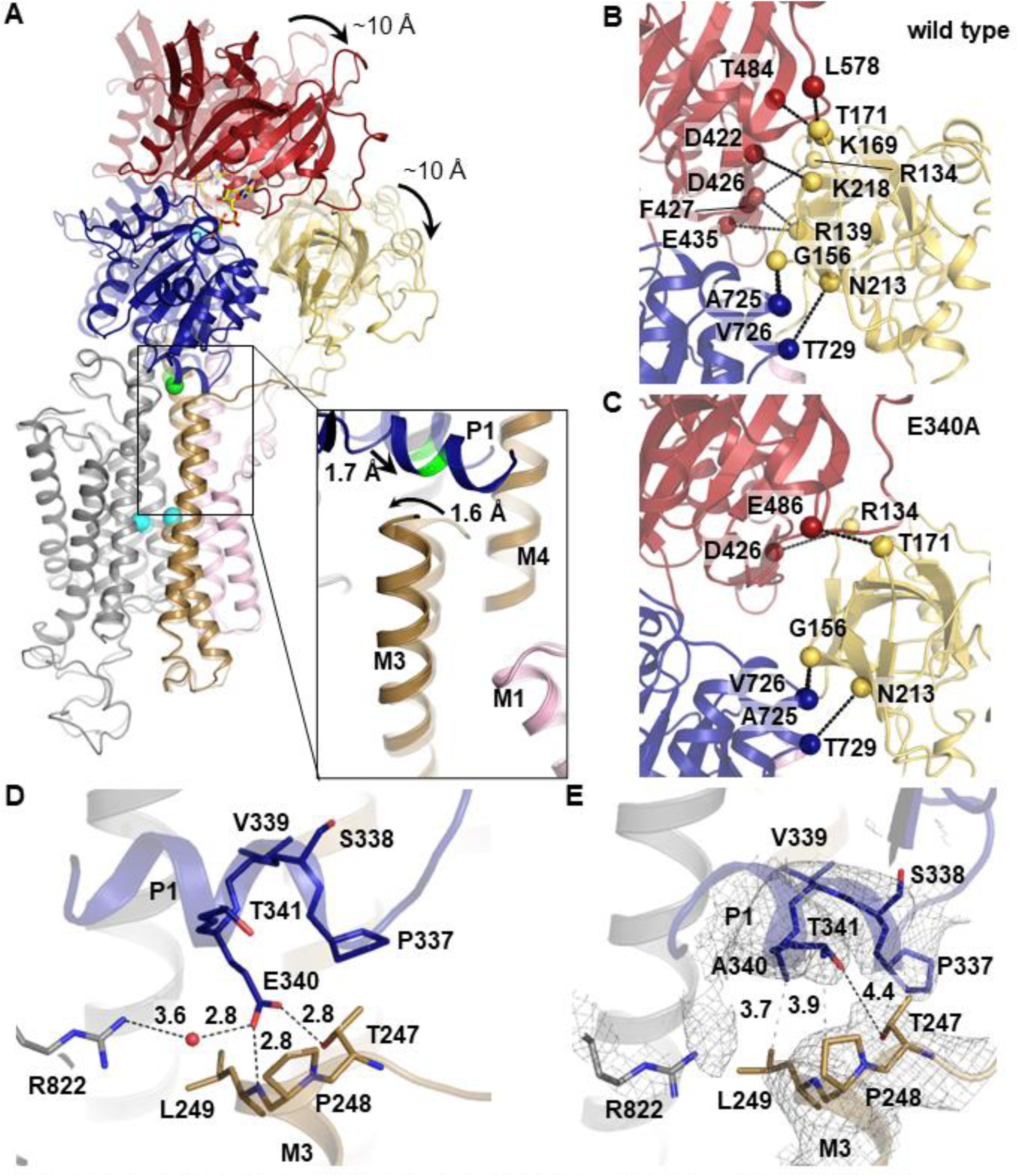
Comparison of SERCA WT (PDB 3N8G) and E340A crystal structures. Colouring is like in Figure 1. A Domain rearrangements in E340A. Superposition of E340A with WT (transparent). Inset: local changes near residue 340. B,C Polar contacts between the A-domain (yellow) and the N-(red) and P- (blue) domains in WT SERCA (B) and E340A (C). D Local hydrogen bond network (black dashes) around residue 340 in WT SERCA. E Local hydrogen bond network (black dashes) and hydrophobic contacts (grey dashes) around residue 340 in E340A. Grey mesh: 2*m*Fo-*D*Fc electron density map contoured at 1.0 σ.

In contrast to the almost identical arrangement in the TM domain, there is a pronounced shift (ca 10 Å) in the position of the cytosolic headpiece, which is rotated further ‘downward’ towards the membrane surface in E340A (Fig 2A and Movie S1). The pivot points of this headpiece movement are at the cytosolic ends of M2, M4 and M5. Whereas M2 and M4 stay very rigid throughout their entire lengths, M5 undergoes a slight bend in the region between its N-terminal end (at Phe740), which is embedded in the P-domain, up till residue Gly750, which sits at the ‘rear side’ of P1, exactly opposite residue 340. On the ‘front side’ of P1, the N-terminal end of M3 moves in the opposite direction of the shifting P1 helix, bending slightly inwards with its first two N-terminal windings. (Fig 2A insert, Movie S1).

Intriguingly, the downward rotation of the cytosolic headpiece in E340A is an extension of exactly the same trajectory as the movement of WT SERCA when it shifts from empty Mg*E*1 to the Ca^2+^-bound *E*1 form (Ca_2_*E*1): when Ca^2+^ binds to WT SERCA, the entire P1-M3 network moves upwards relative to L6-7 and M5, by about 4.5 Å, while the overall headpiece rotates down towards the membrane surface (8). In E340A, this headpiece downward rotation continues further by about 10 Å (as visualized in a superposed morph between the Mg*E*1 (PDB 4H1W) and the Ca_2_*E*1 WT (PDB 3N8G) and E340A (PDB 6RB2) structures, see Movie S2). This extension of the WT Ca^2+^-binding trajectory is reflected in a larger main chain RMSD between the empty WT structure and the Ca^2+^-bound states in E340A (5.6 Å) than in WT SERCA (4.7 Å).

The observation of an ‘overshooting’ Ca^2+^-binding movement in E340A led us to ask whether this mutation might lead to a stabilization of a Ca^2+^ occluded state, the state that follows Ca^2+^- (and nucleotide) binding in the catalytic cycle. Ca^2+^ occlusion is normally stabilized by the initiation of phosphorylation by ATP and not by Ca^2+^ binding alone (7). We hence looked at further structural deviations of SERCA E340A from WT, starting at the contacts between the three domains of the cytosolic headpiece (N, P, and A domains).

### Connectivity between the N- and A domains is looser in E340A

While the domain bodies themselves do not change between WT and E340A (main chain RMSD of 0.37 Å, 0.37 Å and 0.32 Å for the single P-, N-, and A domains, respectively), their concerted movement relative to the rest of the structure in E340A is reflected by an overall main chain RMSD for the entire headpiece of 1.16 Å. The P- and the N domain appear to move together as one rigid body (main chain RMSD = 0.47 Å for the P/N body), whereas the A domain is displaced slightly further, with the distance between the N- and the A domain increasing by approx. 3 Å in the final position. The main chain RMSD of 1.4 Å for the N/A body is the largest RMS deviation measured for any element of the two structures, indicating that this is the largest structural change induced by the mutation. This change leads to the loss of 6 out of 8 polar contacts between the N- and the A domain (Arg134 to Phe427, Arg139 to Asp426 and Glu435, Lys169 to Thr484, Thr171 to Leu578, and Lys218 to Asp422), leaving only two contacts intact between these two domains (Arg134 to Asp426 and Thr171 to Glu486), whereas the A domain’s contacts to the P domain remain intact (Gly156 to Ala725 and Val726, Asn213 to Thr729) (Fig 2B and C). This loss of N-A contacts is also evident from our MD simulations (see below and Fig 3C).

**Figure 3:**
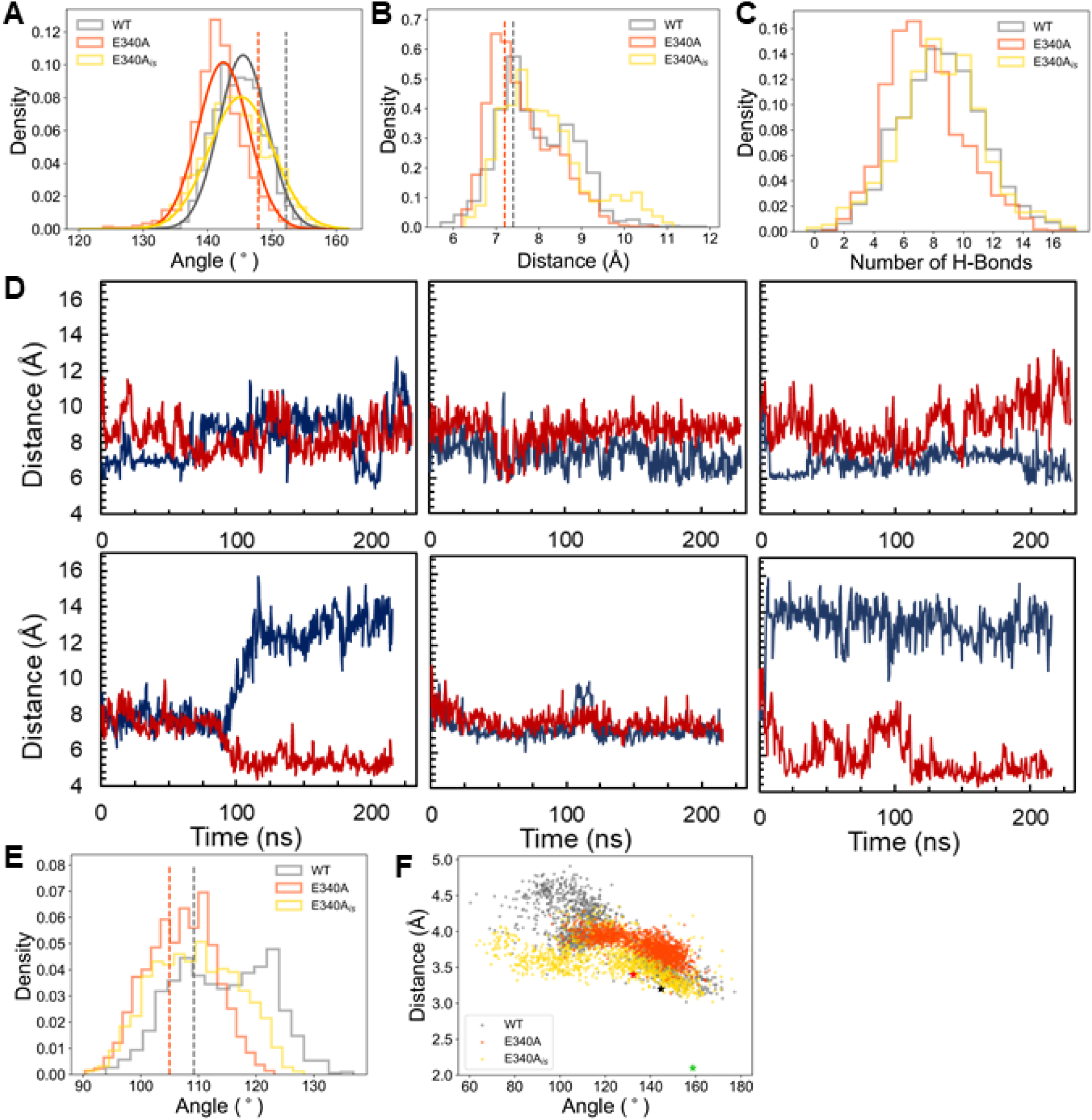
Molecular dynamics simulations of SERCA WT (PDB 3N8G), E340A and modelled mutant E340A_*is*_. WT is shown in gray, E340A in orange and E340A_*is*_ in yellow. Histograms give density as an arbitrary unit on the y-axis. Dotted lines and asterisks refer to reference values in respective crystal structures. A Angle histogram between the soluble headpiece and the transmembrane domain, measured between Leu13 in the A-domain and Thr86 and Leu98 in M2. B Distance histogram between Cα atoms of Leu249 at the tip of M3 and residue 340. C Histogram of hydrogen bonds between the A-domain and the N/P domain body. D Distance traces between Arg833-Nε and Glu/Ala340-Cα (blue) and between Leu249-Cα and Pro824-Cα (red). Left to right: simulation runs 1- 3. Top traces: WT, bottom traces E340A. E Histogram of kink angles in M1, measured between Cα atoms of Trp50, Arg63 and Val74. F Geometry at the catalytic site. Angles between the O- and Pγ-atoms of the terminal phosphoanhydride bond of ATP and the closest carboxyl oxygen (Oδ) of Asp351 versus the distance between Oδ and Pγ. Green asterisk: 1T5T, ADP-AlFx complex.

### Local changes at the mutated residue 340

In the Ca_2_*E*1 form of WT SERCA (PDB 3N8G), the Glu340 side chain is in hydrogen bonding distance to the side chain OH group of Thr247 and the main chain nitrogen of Leu249 at the N-terminal end of M3, with additional electrostatic attraction by the partial positive charge of the helix N terminus. A third interaction of Glu340 is a water-mediated contact to Arg822 in L6-7 (Fig 2D).

Interestingly, the local conformation around the Glu → Ala mutation (within a radius of ∼ 10 Å) is very similar to WT, with measurable shifts of under 2 Å (Fig 2A and D-E). While the local similarity may seem surprising at first glance, looking at the larger global structural changes described above, it becomes clear that residue A340 is located very close to the pivot point for the described headpiece rotation and therefore does not move much itself.

The overall distance between P1 and M3 is slightly smaller than in WT, and the nature of the contact between these two structural elements is very different in E340A: there are no apparent polar interactions between P1 and M3, except for a distant (4.4 Å) contact between Thr341 in P1 and Thr247 in M3 (Fig 2E). Instead, the electrostatic attraction to the partially positive M3 N terminus is replaced with hydrophobic interactions (Ala340 to Leu249, Pro337 to the methyl group of Thr247, and Thr341 to Pro248). In fact, this interaction appears to be rather tight, as shown by our MD simulations (see below and Fig 3B).

Removal of the Glu340 side chain abolishes the contact to Arg822, and the entire L6-7 loop (Phe809 to Ser830) is displaced laterally by ca. 3 Å, increasing the distance between Arg822 and the P1 helix. In order to test the hypothesis that the loss of the Arg822-Glu340 contact in E340A contributes to the functional impairment of this mutation, and to validate the structural and functional differences between WT and E340A described above, we conducted molecular dynamics (MD) simulations.

### Molecular Dynamics Simulations support a more occluded state of E340A

In order to assess whether the differences we found between the crystal structures of WT SERCA and E340A might reflect stable conformations occurring in the native membrane environment, and to find further explanations for the observed functional impairment of the mutant, we carried out 3 × 200 ns of all-atom MD simulations of each structure reconstituted in a POPC lipid bilayer. As an internal control, we generated an *in silico* mutant E340A_*is*_ by truncation of the Glu340 side chain from the WT Ca_2_*E*1 structure and subjected it to the same simulations.

To quantify the conformational downward shift of the cytosolic headpiece relative to the membrane, we determined the angle between three points; Leu13 in the A domain and Thr86 and Leu98 in M2 (Fig S1A). When comparing the two crystal structures, the angle in E340A is 5° more acute (147°) than in WT (152°). During the course of the simulations, both structures relax towards more acute angles, but the difference between them stays about the same, with E340A having a median angle of 142° and WT of 146° (Fig 3A). This confirms that the difference in headpiece angle is a true feature of E340A that persists in a dynamic membrane environment.

Coherent with the above finding, the P1 helix remains closer to the tip of M3 in E340A, as shown by the distance between the C_α_-atoms of residue 340 and Leu249 at the tip of M3 (Figs 3B and S1B). The WT structure exhibits a bimodal distribution of distances with peaks at 7.5 Å and 8.8 Å, whereas E340A has a prominent peak at a shorter distance than WT (6.9 Å).

The loss of connectivity between the A domain and the N/P-body described above is also apparent from the MD simulations. The number of H-bonds between these elements remains reduced in E340A throughout the simulation (Fig 3C).

In the three analyses described above, the *in silico* mutant E340A_*is*_ gives similar distributions to WT, likely because the rather large domain movements are too slow to be seen after the short simulation time of 200 ns. When looking at more local dynamics, however, E340A_*is*_ does behave more similar to E340A, as seen in the analyses below.

The crystal structure suggests that Glu340 can engage in up to three hydrogen bonds: with Thr247, the backbone of Leu249 and a water-mediated contact to Arg822 (Fig 2D). The link to Arg822 in L6-7 is completely lost in E340A (Fig 2E). Despite the loss of this ionic contact, there is only a slight displacement of L6-7, which is held in place by contacts to L8-9 and other regions of the P domain. Accordingly, we could not detect an increased motility of L6-7 in our MD simulations (Fig S2A). Nevertheless, our simulation data show that – in the absence of the Glu340 side chain – Arg822 can swing out of the site between P1 and M3 (Fig S2B). This is measurable as a sudden increase in the distance between Arg822 N_ε_ and Ala340 C_β_ from about 7 Å to 13 Å in E340A and to more than 14 Å in E340A*is* (Figs 3D and S2C). In WT SERCA, Arg822 does not swing out spontaneously, and the distance stays at 7-10 Å. (Fig 3D). Importantly, once moved to the ‘out’ position, Arg822 does not return to the ‘in’ position over the course of any of our simulations of E340A and E340A_*is*_ (Figs 3D and S2C). The outward swing of Arg822 does, however, not lead to a loss of contact between P1, M3 and L6-7, as might be expected at first glance. On the contrary, concomitant with the outward swing of Arg822 from the wedge between P1 and M3, the space is taken up by Leu249 at the tip of M3, which closes in on the gap and approaches Pro824 in the L6-7 loop (Fig 3B and S2D), reducing the distance between the two Cα atoms from 10 Å (E340A*is*) or 7.5 Å (E340A) to 5 Å (Figs 3D and S3C). This approach of M3 toward L6-7 is mutually exclusive with the ‘in’ position of Arg822, as seen by a conspicuous correlation of the distance plots between Arg822 and Ala340 and between Leu249 and Pro824. This correlation is also evident in the *in silico* mutant E340A*is* but absent from the WT (Figs 3D and S3C), suggesting that the changes in the contact network around P1 play a central role in mediating the observed structural changes in E340A.

Comparing the dynamics of the transmembrane segments throughout the simulations, we found a remarkable difference in the kinked region of M1 between WT and both E340A and E340A_is_: M1 is rigidly kinked in E340A, while in WT the kink region displays more dynamics and can straighten to some extent (Figs 3E and S3). This finding is particularly interesting in light of prior studies showing an involvement of M1 with Ca^2+^ binding and occlusion(7, 8).

In order to further probe our simulation data with respect to Ca^2+^ occlusion in E340A, we analyzed the nucleotide binding site. Ca^2+^ occlusion is coupled to phosphorylation of Asp351, and a more occluded conformation of SERCA would be expected to be more prone to be phosphorylated. We hence compared the distances between the ATP γ-phosphate and the respective closer of the two carboxyl oxygen atoms of Asp351, along with the angles between the carboxyl oxygen and the O-P bond of the terminal phosphoanhydride in ATP (Fig S4). For comparison, the crystal structure in which the fully occluded transition state of phosphorylation has been trapped with ADP-AlF_x_ (7) the distance and angle between the Asp351 O_δ_ and the aluminum atom are 2.1 Å and 159°, respectively. As seen in Fig 3F, the WT structure displays a large spread of angles (between 60 and 180°) and distances (between 3 and 5 Å), many of which are not compatible with an in-line associative hydrolysis mechanism. In E340A, the distances take up a much narrower spread mostly between 3.5 and 4 Å, and the angles are more restricted towards the catalytically competent obtuse region (between 110° and 170°). E340A*is* displays an intermediate distribution. Our data are therefore consistent with a more occluded state in both the ion binding and the nucleotide binding regions of E340A.

## Discussion

We have addressed the molecular basis for intramolecular communication in SERCA by analyzing the structure and functional properties of the SERCA mutant E340A. The Glu340 residue is strongly conserved among P-type ATPase membrane transporters and is located at a central position of the ATPase, at the interface between the phosphorylation domain and transmembrane segment, where the Ca^2+^ binding sites are located.

The structure of the mutant differs from the equivalent wild-type structure in: i) a lowered cytosolic headpiece relative to the membrane ii) altered connectivity between the nucleotide-binding and the actuator domain, iii) a shift in the Ca^2+^-gating transmembrane helix M3. MD confirms the structural changes observed and moreover reveals altered dynamics in regions involved in Ca^2+^-occlusion, in particular a rigidly kinked M1. These structural findings point to a stabilization of a more occluded conformation and are discussed below in relation to the functional properties of the E340A mutant.

The downward movement of the cytosolic headpiece towards the membrane in E340A is an extension of exactly the same trajectory as the movement of WT SERCA when it shifts from empty Mg*E*1 to the Ca^2+^-bound *E*1 form (Ca_2_*E*1). Hence, the loss of Glu340 allows an ‘overshoot’ of this closure movement after ion binding, which is pivoting around P1. At the same time, within the headpiece, the A domain gets displaced and loses more than half of its contacts with the N domain. The exact functional implication of this effect is not entirely clear. The A domain undergoes by far the largest movements during the catalytic cycle, and through its direct linkage to TM helices M1-M3 it directly affects the geometry of the Ca^2+^ binding sites. On the other hand, its position relative to the N- and P domains dictates whether the site of ATP hydrolysis can adopt a catalytically competent conformation or not. Hence, any alteration of the dynamics of the A domain will coactively alter SERCA catalysis.

The striking similarity of the Ca^2+^ binding sites and their immediate surroundings in the WT and E340A crystal structures indicates that the differences in the Ca^2+^ binding and dissociation behavior are not caused by a perturbation of the geometry at the Ca^2+^ sites in the TM domain but rather by altered kinetics of the actual binding process. One of the few structural differences in the transmembrane region is a small inward shift of the tip of M3 in E340A. Interestingly, a mutation of Leu249 at the tip of M3 to alanine leads to increased rates of both Ca^2+^ binding and dissociation (21). Furthermore, in the SERCA Ca^2+^ binding site mutant E309Q, which is defective in both Ca^2+^-binding and occlusion, the tip of M3 is outward-shifted (19), highlighting the immediate influence of small changes to the spacing between P1 and M3 for the ion binding dynamics in SERCA.

The loss of the electrostatic interaction between Glu340 and Arg822 in L6-7 appears to have only a small structural effect at first glance. However, prior studies have implicated an important role of L6-7 in SERCA catalysis (15, 16, 21). For example, the D813A/D818A mutation leads to a dramatic loss of Ca^2+^ affinity (15). There are also conspicuous similarities between the behavior of the mutant E340A and some L6-7 mutants, including R822A: a slowing of the Ca^2+^-binding transition from *E*2 and of Ca^2+^ dissociation (16). The loop L6-7 has moreover been suggested to be part of an ion access pathway in both SERCA and the Na^+^/K^+^-ATPase (11, 12, 14, 15, 22), and/or to contribute to the coordination of events between the cytosolic and transmembrane domains (13, 15, 16). In light of our finding that the E340A mutation allows for an irreversible swinging out of Arg822, which is correlated to a concomitant inward movement of M3, the reverse conclusion must be that the interaction network between Arg822, Glu340 and Leu249 is critical for maintaining a proper architecture of the Ca^2+^ entry and exit pathways.

Finally the ‘rigidification’ of the M1 kink which we see in our MD simulations is also in line with a shift of E340A towards a slightly more occluded state than WT. The kink at Leu60 in M1 permits the conserved residue Phe57 to engage in a hydrophobic cluster that shields off the Ca^2+^ binding sites as a prerequisite for ion occlusion. Notably, a straight M1 helix has been seen in nucleotide-free inward-open E1 structures (19, 23), and is associated with an un-occluded Ca_2_*E*1 state of SERCA. Most notably, when probing the reported interrelation of ion occlusion and phosphorylation (7), we find the geometry of the phosphorylation site to be more narrowly clustered around catalytically competent values.

### How does the structural stabilization of the occluded state manifest itself in functional terms?

Our tryptophan fluorescence data obtained with the detergent solubilized yeast enzyme used for crystallization show that the E340A mutation caused a pronounced slowing of the reaction sequence H_n_*E*2 -> *E*1 -> Ca_2_*E*1 consisting of proton release, Ca^2+^ binding, and the associated conformational changes). This effect explains the reduced ATPase activity of the E340A mutant relative to WT, and the slowing agrees with the change in the dynamics of Ca^2+^ gating suggested from the structural changes in relation to the Ca^2+^-binding sites.

The E340A mutant has previously been analyzed functionally following expression in COS cells (16), showing effects qualitatively similar to the present results. In accordance with our finding of a more occluded E340A structure, the previous functional analysis in COS cell microsomes (16) showed that the E340A mutation not only slows down Ca^2+^ binding, but also the reverse reaction Ca_2_*E*1 -> *E*1 (Ca^2+^ dissociation from the high affinity sites towards the cytosol). The mutation was moreover found not to affect SERCA’s apparent vanadate affinity (which is typically affected by *E*2-*E*1 shifts), suggesting that the slowing of H_n_*E*2 -> *E*1 -> Ca_2_*E*1 is due to inhibition of the latter part of this reaction sequence (*i*.*e*. actual Ca^2+^ binding). The mutation also had no significant effect on the remaining partial reaction steps of the cycle (16). Hence, the kinetic constraints observed for the E340A mutant are consistent with the hypothesis that perturbation of the hydrogen-bonding network around Glu340 causes a slowing, in both directions, of the structural changes associated with the binding of the second Ca^2+^ ion to the Ca*E*1 state and the subsequent occlusion step.

In conclusion, our data suggest that E340A represents a structure that is stabilized in a state closer to occlusion and phosphorylation than the WT structure, hence slowing Ca^2+^ entry and release but favoring phosphorylation by ATP. Conversely it can be concluded that the conserved residue Glu340 in SERCA has an important role in maintaining the structural flexibility needed to allow for rapid Ca^2+^ exchange at the membranous binding sites, and in relaying this flexibility to the site of phosphorylation.

This key function provides an explanation for the evolutionary acquisition and strict conservation of this glutamate residue throughout the entire P-type ATPase family, irrespective of their ion specificity.

## Materials and Methods

### Chemicals

Octaethylene glycol mono-n-dodecyl ether (C_12_E_8_) was purchased from Nikkol Chemical (#BL-8SY; Tokyo, Japan), and n-dodecyl β-D-maltopyranoside (DDM) was from Anatrace (#D310; Maumee, OH). Streptavidin Sepharose high performance resin was provided by GE Healthcare (#17-5113-01; Orsay, France). Thapsigargin (TG stock solution was prepared at 1 mg/mL in DMSO, i.e. about 1.5 mM) was from VWR International (#586005; Fontenay-sous-Bois, France). All other chemical products were purchased from Sigma (Saint-Quentin Fallavier, France). Sequence for SERCA1a heterologous expression was from rabbit.

### Cloning, expression and purification

SERCA1a E340A cDNA (SERCA1a[E340A]) was recovered from pMT2 vector initially used for expression in COS cells (16) and cloned into the yeast expression plasmid pYeDP60, with a C-terminal biotin acceptor domain (24), resulting in pYeDP60_SERCA1a[E340A]-Bad. The construct was checked by sequencing (Eurofins MWG). The *Saccharomyces cerevisiae* yeast strain W303.1b/Gal4 (α, leu2, his3, trp1::TRP1-GAL10-GAL4, ura3, ade2–1, canr, cir) was the same as previously described (24). Transformation was performed according to the lithium acetate/single-stranded carrier DNA/PEG method (25).

Expression and purification of the E340A mutant SERCA1a was done as previously described for the WT (WT) enzyme (24). Briefly, after overexpression in the yeast *S. cerevisiae*, light membranes were prepared and solubilized by DDM for subsequent purification by Streptavidin affinity chromatography (24, 26, 27). Purified WT SERCA1a was recovered in a buffer containing 50 mM MOPS-Tris pH 7, 100 mM KCl, 5 mM MgCl_2_, 2.1 mM CaCl_2_, 40 % glycerol (v/v) and 0.5 mg/mL DDM, together with some thrombin remaining from the elution procedure. The protein concentration in the purified fraction was typically in the 0.05 to 0.15 mg/mL range depending on the batch. Further purification for crystallization trials was carried out as described previously for the SERCA E309Q mutant (19), involving exchange of DDM with C_12_E_8_ and relipidation of the purified SERCA with DOPC.

### ATPase activity measurements

Steady-state ATPase activity measurements were performed using an enzyme-coupled assay by measuring the rate of NADH oxidation (followed by absorbance at 340 nm) in the presence of 0.02 mg/mL lactate dehydrogenase, 0.04 mg/mL pyruvate kinase, 1 mM phosphoenolpyruvate, 0.3 – 0.4 mM NADH, 5 mM MgATP (28) and in the additional presence of 1 mg/mL C_12_E_8_ and 50 µM free Ca^2+^ to limit time-dependent inactivation of the SERCA1a (29), at 30°C and pH 7.5.

### Tryptophan fluorescence

Intrinsic fluorescence was measured with a Fluorolog spectrofluorimeter (Horiba / Jobin-Yvon, Palaiseau, France), in a temperature-regulated and continuously stirred 2 mL quartz cuvette. Excitation and emission wavelengths were set at 295 and 320 nm, with bandwith of 2 and 5 nm, respectively. Integration time for recording of the signal was 2 seconds. SERCA1a intrinsic fluorescence changes were measured with purified WT or purified E340A mutant suspended at a protein concentration of about 10 µg/mL in a buffer containing 50 mM Mes-Tris pH 6.5, 5 mM MgCl_2_, 20% glycerol (v/v) and 2 mg/mL DDM, at 20°C. Initial Ca^2+^ concentration was adjusted to 105 µM on top of the contaminating Ca^2+^ (3 - 5µM) already present in the buffer. This was followed by the addition of 5 mM EGTA (EG), reducing [Ca^2+^]_free_ to about 100 nM. An extra addition of 12.5 mM CaCl_2_ (Ca) allowed to reach a final [Ca^2+^]_free_ of about 7.6 mM. to recover the initial level of fluorescence. Data were fitted with the GraphPad Prism program with an exponential two phase association law on the first 60 seconds of the raw data following Ca^2+^ addition.

### Crystallization

Purified SERCA E340A at a concentration of 8-10 mg/ml was supplemented with Ca^2+^, Mg^2+^, and AMPPCP to 10, 3, and 1 mM, respectively, and crystallization trials were carried out using the vapour diffusion technique. The crystallization drops were prepared as 1 µl sitting drops, with each drop containing a 1:1 mixture of the concentrated, DOPC-relipidated, and C_12_E_8_-solubilized Ca^2+^-ATPase solution and the reservoir solution. Diffraction-quality crystals were obtained with reservoir solutions containing 200 mM lithium acetate, 18-22% PEG6K, 6-9% glycerol, and 3% tert-butanol. The E340A crystals varied markedly in appearance from crystals obtained with WT under similar conditions. Hence, the E340A crystals displayed a rectangular stick morphology, whereas WT crystals were diamond shaped (cf. Fig S2 in (19)).

### Data collection, processing and refinement

Crystals were flash frozen and tested at the microfocus beam line ID23-2 at ESRF. The best dataset diffracted to 3.2 Å. In accordance with the variant crystals form, also the space group and unit cell parameters were different from WT SERCA. Hence, WT crystallizes in space group C2 with unit cell parameters (163 Å, 76 Å, 151 Å, 90°, 109°, 90°), and E340A in space group *P*2_1_2_1_2 with unit cell parameters (126 Å, 232 Å, 50 Å, 90°, 90°, 90°), which are hitherto unknown parameters for SERCA. Phasing was done by molecular replacement with the Ca_2_*E*1-AMPPCP form (1T5S). Initial rigid-body refinement with each domain as a single rigid body was followed by all-atom and TLS refinement in *PHENIX* (30).

Molecular graphics figures were prepared with PyMOL (31) and morphs with Morphinator (32).

### Molecular Dynamics Simulations

Simulations were built using the coordinates from PDB 3N8G (WT) and 6RB2 (E340A). The protein atoms were described using the CHARMM36 force field (33), and were built into a lipid bilayer composed of POPC molecules and solvated with TIP3P water and Na^+^ and Cl^-^ to 150 mM in a 120 × 120 x 180 Å box. Systems were built using CHARMM-GUI (34, 35). Generation of the *in silico* mutant E340A_*is*_, was done with CHARMM-GUI.

The systems were energy minimized using the steepest descent method, then equilibrated with positional restraints on heavy atoms for 100 ps in the NPT ensemble at 310 K with the V-rescale thermostat and semi-isotropic Parrinello-Rahman pressure coupling (36, 37). Production simulations were run in triplicate without positional restraints, with 2 fs time steps for a minimum of 200 ns. All simulations were run using GROMACS 2019 (38). Simulations were analysed using GROMACS tools and images were made using VMD (39). Graphs were plotted using Matplotlib (40).

## Acknowledgments

This work was supported by the French Infrastructure for Integrated Structural Biology (FRISBI; ANR-10-INSB-05), and the Centre National de la Recherche Scientifique (CNRS), the Agence Nationale de la Recherche and the Domaines d’Intérêt Majeur Maladies Infectieuses Région Ile de France (to MlM, CJ, GL and CM)), the Center for membrane Pumps in Cells and Disease (PUMPkin) of the Danish National Research Foundation (to JVM, PN, JPA and MB), the Lundbeck Foundation (to PN), the Wellcome Trust (ref. 220063/Z/20/Z to MMGG), and the Danish Council for Independent Research (to JPA). We are grateful to Anna Marie Nielsen for technical support and Jesper L. Karlsen and Mark Sansom for scientific computing facilities.

