## Supplementary Information for "The SERCA residue Glu340 mediates inter-domain communication that guides Ca^2+^ transport"

**This PDF file includes:**

Figures S1 to S5
Tables S1 to S2
Legends for Movies S1 to S2

**Other supplementary materials for this manuscript include the following:**

Movies S1 to S2

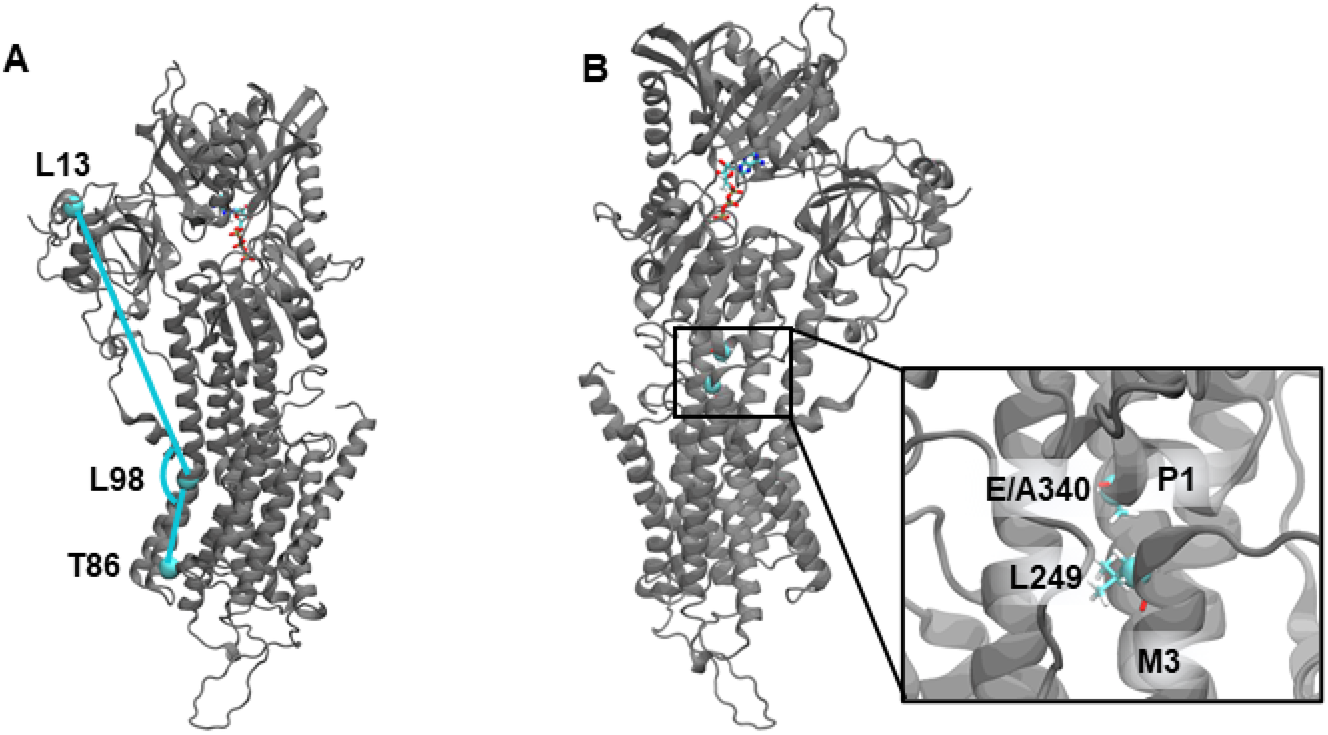


**Figure S1: Visualisation of MD simulation analyses of E340A conformation**

A. Residues used to determine the headpiece angle, Leu13, Thr86 and Leu98.
B. Location of residues Glu/Ala 340 on P1 and Leu249 in M3

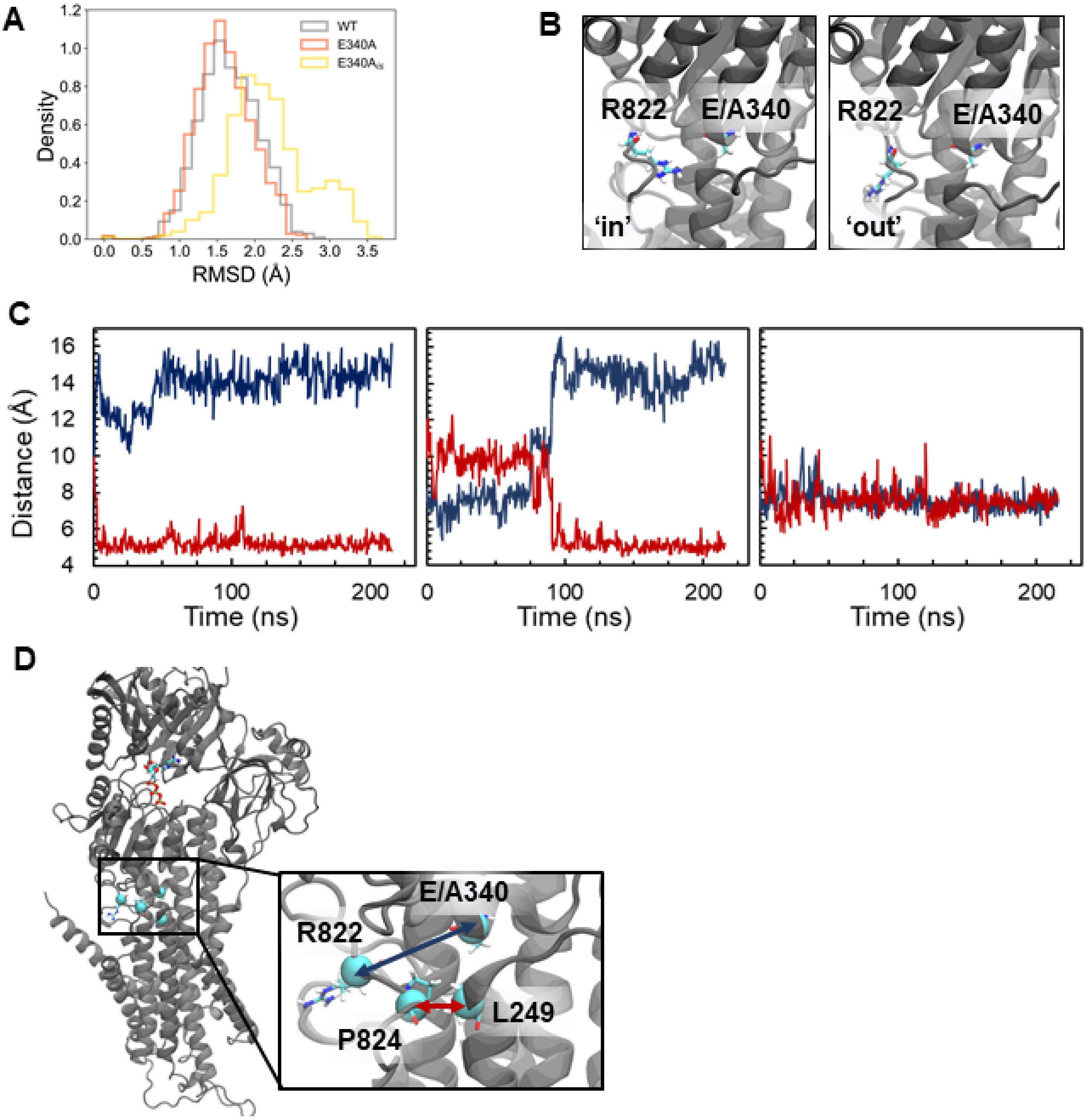


**Figure S2: MD simulation analysis of the L6-7 loop.**

A	RMSD histogram of L6-7 over the course of the simulation.
B	'In' (left) and 'out' (right) positions of Arg822.
C	Distance traces between Arg833-Nε and Glu/Ala340-$C_\alpha$ (blue) and between Leu249-$C_\alpha$ and Pro824-$C_\alpha$ (red) in $E340A_{ls}$. Left to right: simulation runs 1-3.
D	Visualisation of the distance measurements between Glu/Ala 340 and Arg822 and Leu 249 and Pro824.

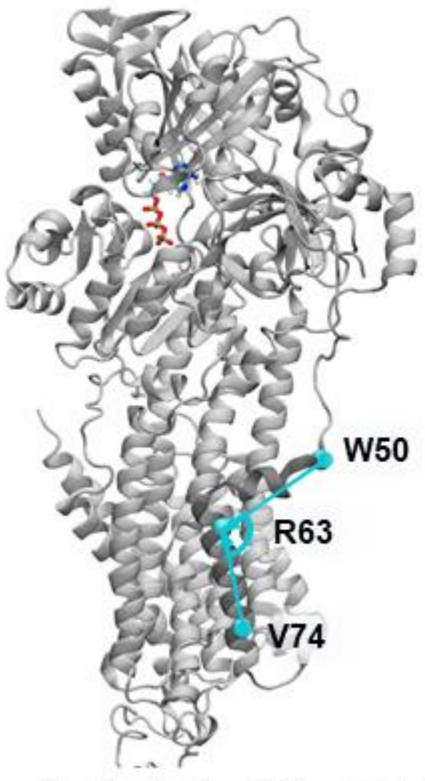


Figure S3: Visualisation of MD analysis of the M1 kink, measured between residues Trp50, Arg63 and Val74.

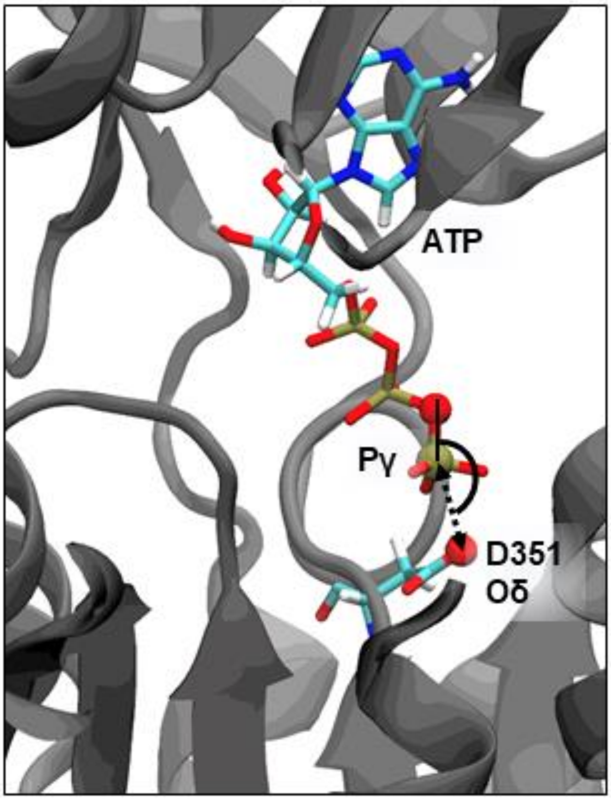


**Figure S4: Visualisation of MD analysis of the phosphorylation site geometry.**

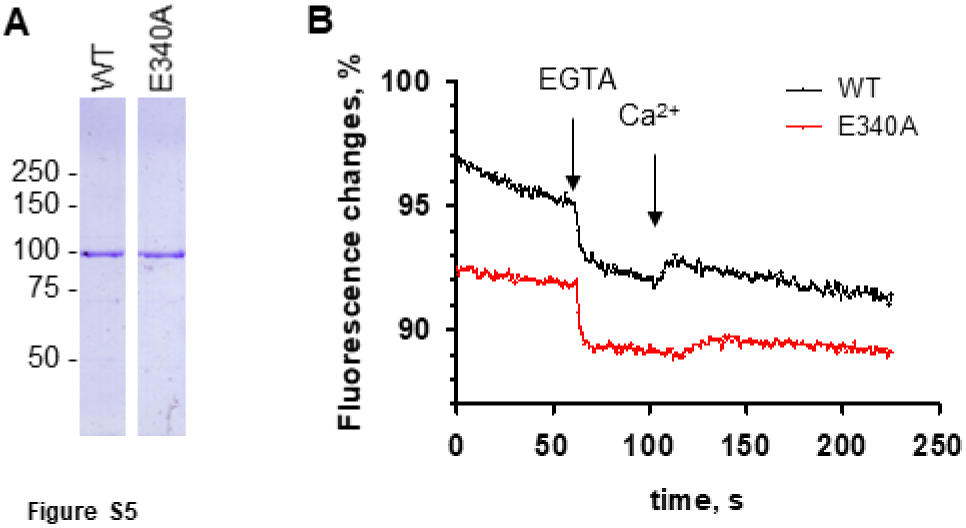


**Figure S5**

A Coomassie blue stained SDS-PAGE of the purified SERCA WT (WT) and E340A proteins.

B SERCA1a intrinsic fluorescence changes measured with purified WT (black) or E340A (red) starting at 105 μM $Ca^{2+}$ followed by addition of 5 mM EGTA and 12.5 mM extra $Ca^{2+}$. Fluorescence intensities are given as a percentage of the fluorescence level at start. For sake of clarity, data for E340A were shifted down manually by 10%. The traces shown are the average of 4 experiments for WT and 3 experiments for E340A.

**Table S1:** SERCA Glu340 is conserved in ion- and lipid transporting P-type ATPases.

In the P-type ATPase database (http://www.traplabs.dk/patbase/Motifs.html), the glutamate is conserved in 135 out of 159 P-type ATPases. The P3B and P5 subfamilies are the only groups where it systematically misses, being replaced by N and Q. In P1B, a few bacterial pumps have Q, S, or D.

| **P-type Family** | **Protein** | **Substrate** | **Organism** | **P1 Sequence** |
|---|---|---|---|---|
| P1A | KdpB | $K^+$ | *E. coli* | RAVEAAGD |
| P1B | ATP7A | $Cu^+$ | *H. sapiens* | EPLEMAHK |
| P2A | SERCA1A | $Ca^{2+}$ | *H. sapiens* | PSVETLGC |
| **P2A** | **SERCA1A** | **$Ca^{2+}$** | ***O. cuniculus*** | PSVETLGC |
| P2B | PMCA | $Ca^{2+}$ | *H. sapiens* | DACETMGN |
| P2C | $Na^+$,$K^+$-ATPase | $Na^+$, $K^+$ | *H. sapiens* | EAVETLGS |
| P2C | $H^+$,$K^+$-ATPase | $H^+$, $K^+$ | *H. sapiens* | EAVETLGS |
| P2D | CTA3 | $Ca^{2+}$ | *S. pombe* | EALEALGG |
| P3A | AHA2 | $H^+$ | *A. thaliana* | TAIEEMAG |
| P3A | Pma1 | $H^+$ | *N. crassa* | SAIESLAG |
| P3B | MgtA | $Mg^{2+}$ | *E. coli* | DAIQNFGA |
| P4 | ATP8B1 | Phospholipid | *H. sapiens* | TLNEQLGQ |
| P5 | ATP13A2 | Polyamine | *H. sapiens* | QRINVCGQ |

**Table S2:** Crystallographic data collections and refinement statistics. Values in parentheses refer to the highest resolution shell

| **Data collection** | |
|---|---|
| Beamline | ESRF ID23-2 |
| Space group | $P2_12_12$ |
| Unit cell (Å, °) | a=232.90, b=126.99, c=49.81, α=β=γ=90 |
| Wavelength (Å) | 0.8729 |
| Resolution (Å) | 75-3.2 (3.3-3.2) |
| Number of unique reflections | 25337 (2201) |
| Completeness (%) | 99.9 (99.8) |
| Multiplicity | 6.1 (6.2) |
| I/σI | 10.9 (1.3) |
| Rmeas | 0.196 (>100) |
| $CC_{1/2}$ in highest resolution shell | 0.47 |
| Wilson B-factor | 97.8 |
| **Refinement** | |
| Resolution (Å) | 49.81-3.2 |
| Rwork/Rfree (%) | 0.21/0.26 |
| Rmsd bond (Å) | 0.009 |
| Rmsd angle (°) | 0.691 |
| Mean B-factors ($Å^2$) | 90.7 |
| Ramachandran plot (%) Favoured/allowed/outliers | 93.0/6.4/0.6 |

**Movie S1 (separate file). Morph between equivalent $Ca_2$E1 crystal structures of SERCA WT (PDB 3N8G) and E340A (PDB 6RB2).** The inset is a zoom on the region around residue 340 (green sphere).

**Movie S2 (separate file). Superposed morphs between crystal structures of $Ca^{2+}$-free E1 WT (PDB 4H1W) and $Ca_2$E1 forms of WT (PDB 3N8G, light colors) and E340A (PDB 6RB2, dark colors).** The inset is a zoom on the region around residue 340 (green sphere).
